# Phylogeny, phylogeography and hybridization of Caucasian barbels of the genus *Barbus* (Actinopterygii, Cyprinidae)

**DOI:** 10.1101/473173

**Authors:** Boris A. Levin, Alexander A. Gandlin, Evgeniy S. Simonov, Marina A. Levina, Anna E. Barmintseva, Bella Japoshvili, Nikolai S. Mugue, Levan Mumladze, Namig J. Mustafayev, Andrey N. Pashkov, Haikaz R. Roubenyan, Maxim I. Shapovalov, Ignacio Doadrio

## Abstract

The phylogenetic relationships and the phylogeography of seven species of Caucasian barbels of the genus *Barbus* s. str. were studied based on extended geographic coverage and the use of mtDNA and nDNA markers. Based on the 26 species studied, matrilineal phylogeny of the genus *Barbus* is composed of two clades: a) West European clade, and b) Central and East European clade. The latter comprises two subclades: b1) Balkanian subclade, and b2) Ponto-Caspian subclade, which includes 11 lineages mainly from Black and Caspian Sea drainages. Caucasian barbels are not monophyletic and are subdivided into two groups. The Black Sea group encompasses species from tributaries of the Black Sea, including the reinstalled *B. rionicus*, except for *B. kubanicus*. The Caspian group includes *B. ciscaucasicus, B. cyri* (with *B. goktschaicus*, which might be synonymized with *B. cyri), B. lacerta* from the Tigris-Euphrates basin and *B. kubanicus* from the Kuban basin. The genetic structure of Black Sea barbels was influenced by glaciation-deglaciation periods accompanied by freshwater phases, periods of migration and the colonization of Black Sea tributaries. Intra- and intergeneric hybridization among Caucasian barbines was revealed for the first time. In the present study, we report the discovery of *B. escherichii* in the Kuban basin, where only *B. kubanicus* was known to inhabit. Hybrids of these two species were detected based on both mtDNA and nDNA markers. Remarkably, the Kuban population of *B. escherichii* is distant to closely located conspecific populations, and we consider it as a relic. We reveal the intergeneric hybridization between evolutionary tetraploid (2n=100) *B. goktschaicus* and evolutionary hexaploid (2n=150) *Capoeta sevangi* in Lake Sevan.

## 1. Introduction

Polyploid cyprinids are a large group widely distributed in Eurasia and Africa. The freshwater fishes of the genus *Barbus* s. str. are of a tetraploid lineage (2n=100) of Eurasian barbines (Berrebi and Tsigenopoulos, 2003; Ráb and Collares-Pereira, 1995; Vasiliev, 1985) composed of 35 valid species widely distributed from the Iberian Peninsula in Western Europe to the Transcaspian region (the Atrek basin) in Central Asia (Eschmeyer, 2018). This lineage is phenotypically and genetically distinct from another tetraploid lineage of Eurasian barbels, the genus *Luciobarbus* (Doadrio, 1990; Doadrio et al., 2002; Levin, 2004).

The mitochondrial relationships of the European *Barbus* s. str. and the phylogeography of some species were clarified, mainly in the beginning of the 21st century (Doadrio et al., 2002; Kotlík and Berrebi, 2001, 2002; Kotlík et al., 2004; Tsigenopoulos et al., 2002). Involvement of nuclear markers in this polyploid group promoted further uncovering of the evolutionary patterns of the *Barbus* diversity, an elaboration of the population structure, hybridization, and gene flow in certain groups of European barbels in Iberian (Gante et al., 2009, 2015; Machordom et al., 1990), Balkan (Marková et al., 2010), and Italian (Buonerba et al., 2015; Meraner et al., 2013) peninsulas. Molecular expertise coupled with morphological analysis resulted in the discovery of several new species, which were subsequently described as *Barbus carpathicus* Kotlík, Tsigenopoulos, Ráb and Berrebi, 2002 (Kotlík et al., 2002) and *Barbus biharicus* Antal, László and Kotlík, 2016 (Antal et al., 2016) from the Danube River, as well as some undescribed cryptic entities (Marková et al., 2010).

In contrast to the barbels from Western and Central Europe, Caucasian species appeared sporadically in some phylogenetic studies (Durand et al., 2002; Kotlík et al., 2004; Zardoya and Doadrio, 1999). Caucasus is a large mountain system located between the Black and Caspian seas and is considered to be one of the world’s biodiversity ‘hotspots’ under conservation priority (Myers et al., 2000). Caucasian freshwater fish fauna is rich in endemism (Levin et al., 2018; Naseka, 2010). In fact, every second species in Caucasus ecoregions are endemic, according to a review of Naseka (2010). Five of six Caucasian barbels are considered Caucasian endemics (Berg, 1949; Naseka, 2010).

The Terek barbel *B. ciscaucasicus* Kessler, 1877 inhabits the Terek and Kuma riverine system; it is known from other tributaries of the Caspian Sea in East Ciscaucasia, southward from the Sulak basin to the Khudat R. and Kurakh-chai system in northern Azerbaijan (Abdurakhmanov, 1962; Bogutskaya, 2003a). Recently, the range of this species in East Transcaucasia was expanded ca. 100 km southward to the system of the dead-end Pirsagat R. (Gandlin et al., 2017). The Kura barbel *B. cyri* De Filippi, 1865 inhabits the extended Kura and Araks riverine systems, the Caspian Sea drainage, as well as the southward-located tributaries of the Caspian Sea until the Transcaspian Atrek basin (Bogutskaya, 2003b). The Sevan barbel *B. goktschaicus* Kessler, 1877 dwells in the large, isolated Lake Sevan and its tributaries (Bogutskaya, 2003c; Levin and Rubenyan, 2006). The Kuban barbel *B. kubanicus* Berg, 1912 inhabits the Kuban drainage, which is the extended riverine system of West Ciscaucasia. This species was also detected in the Manych system due to recent water diversion from the Kuban tributaries via canals (Poznyak, 1987); the Manych valley served in the past as a connecting passage between the Caspian and Black Sea basins (Esin et al., 2016). The Rioni barbel *B. rionicus* Kamensky, 1899, was described as originating from the Rioni basin, West Transcaucasia, but was later found to be synonymous with *B. escherichii* Steindachner, 1897 by Berg (1914). Based on recent taxonomic opinion, the Rioni barbel may be a valid species (Bayçelebi et al., 2015; Çiçek et al., 2016). The sixth *Barbus* species, Ankara barbel *B. escherichii*, is non-endemic to Caucasus. This species is distributed from the southwestern tributaries of the Black Sea in Anatolia, Turkey, to the Eastern Black Sea drainage on the Russian coast (Eschmeyer, 2017; Reshetnikov et al., 2003). We have to stress that some other *Barbus* species were described in Caucasus but were later transferred to synonyms of previously recognized species. Among these taxa are the Chorokh barbel *B. artvinica* Kamensky, 1899, the Toporovani barbel *B. toporovanicus* Kamensky, 1899, and the Bortschala barbel *B. bortschalinicus* Kamensky, 1899.

Although some genetic studies included the Caucasian *Barbus* species, they were based on few mitochondrial DNA sequences of the cytochrome *b* (*Cytb*) or cytochrome oxidase I subunit (CÒI) of some species (Durand et al., 2002; Khaefi et al., 2017; Kotlík et al., 2004; Zardoya and Doadrio, 1999), while none of Caucasian taxa were sequenced for nuclear markers. This gap in knowledge of the genetic diversity and phylogeny of the significant part of the genus impedes studies on the phylogenetic organization of the *Barbus* s. str., and the evolutionary history of Caucasian species as well as their taxonomy.

The purposes of the present study were i) to estimate phylogenetic relationships, ii) to clarify phylogeography and iii) to contribute to the taxonomy of the Caucasian barbels of the genus *Barbus* s. str. of all the abovementioned Caucasian taxa with wide geographic coverage from Armenia, Azerbaijan, Georgia, and Russia. We also aimed to verify the hybridization among Caucasian taxa, given that such events were repeatedly reported for the *Barbus* from Central and Southern Europe (Buonerba et al., 2015; Lajbner et al., 2009; Machordom et al., 1990; Meraner et al., 2013). For these purposes, we used the datasets of two mitochondrial markers and one nuclear marker.

## 2. Material and methods

### 2.1. Sampling, DNA extraction, and sequencing

DNA samples were collected from 277 individuals of Caucasian and Crimean *Barbus* with local authority permission. All *Barbus* species of this region were collected from 81 sampling sites of the drainages of the Black and Caspian Seas, including the type localities or type basins, except that for *B. escherichii*, which was described by Steindachner (1897) as originating from the Sakarya River, Turkey (Table S1-S2, Figure 1). The mursa *Luciobarbus mursa* (Güldenstädt, 1773) and the khramulya *Capoeta capoeta* (Güldenstädt, 1773) were also sampled for DNA extraction, PCR and subsequent phylogenetic analysis in the Caucasian rivers (Table S1) as outgroups according to previous phylogenetic studies (Levin et al., 2012; Tsigenopoulos et al., 2003). A fragment of pelvic fin was cut and stored in microtubes in 96 % ethanol and deposited in the IBIW RAS Ichthyological collection. DNA-vouchers from each site were killed with overdoses of MS222 and preserved in 10 % formalin or 96 % ethanol and stored in the IBIW RAS Ichthyological collection.

**Fig. 1.**
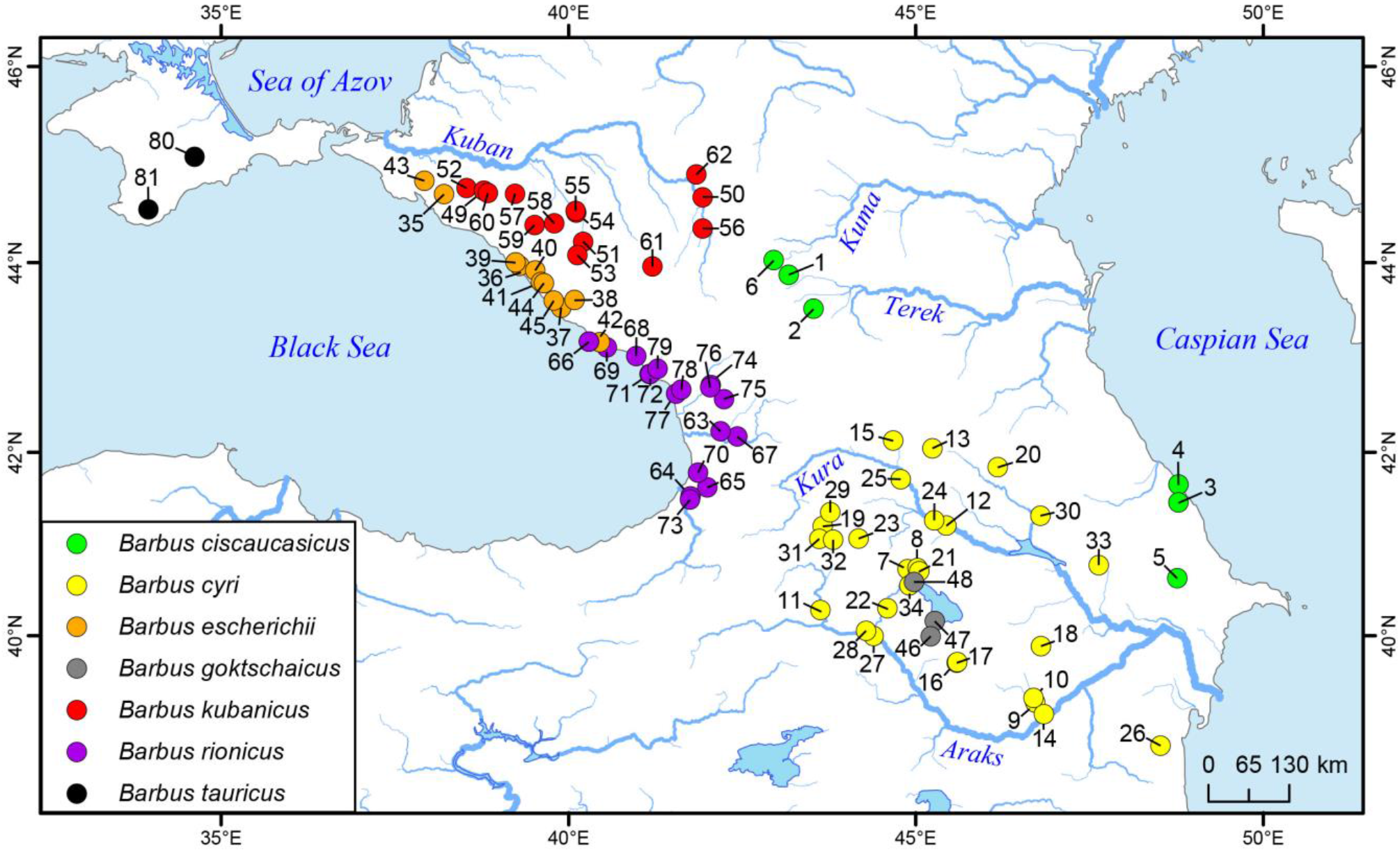
Map of sampling locations. See Table S1 for the corresponding localities numbers.

DNA was extracted using the salt method (Aljanabi and Martinez, 1997). Two mitochondrial markers (barcoding cytochrome oxidase subunit I – *CO*I, and cytochrome *b* - *Cytb*) and one nuclear marker (second intron of beta actin gene – *Act-2*) were amplified using specific primers (Table 2). Barbels are tetraploid with a chromosome number of 2n=100, so that the primers were designed to selectively amplify a single paralogous copy of *Act-2* (Marková et al., 2010). The PCR conditions followed protocols from Ivanova et al. (2007) for *CO*I, Levin et al. (2012) for *Cytb*, and Marková et al. (2010) for the *Act-2* second intron.

**Table 1.**
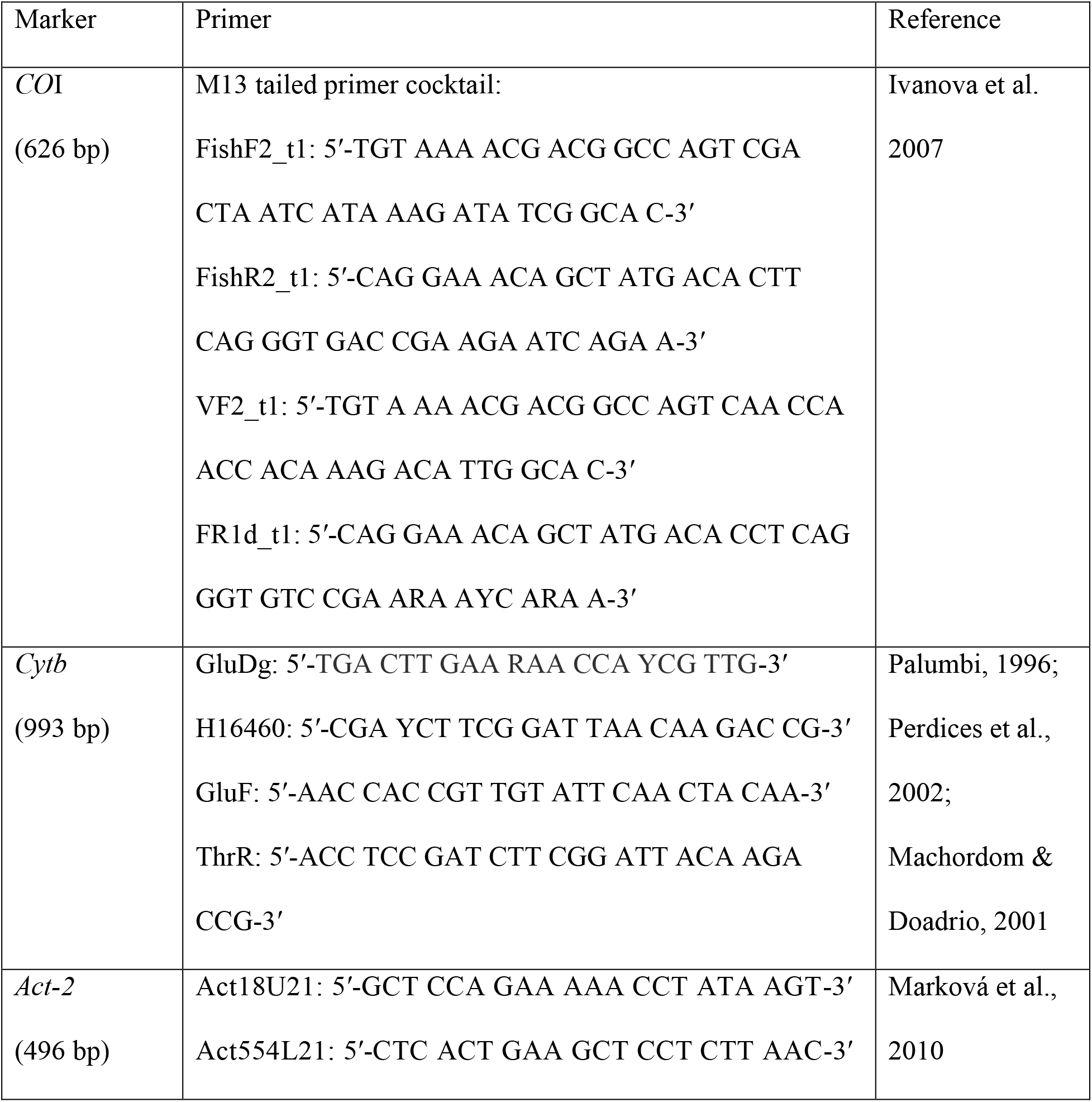
Markers used and primer sequences.

Sequencing of the PCR products purified by ethanol and ammonium acetate (3 M) precipitation was conducted using the Applied Biosystems 3500 DNA sequencer with forward sequencing primer M13F 5’-GTA AAA CGA CGG CCA GT-3’ and reverse sequencing primer M13R-pUC 5’-CAG GAA ACA GC T ATG AC-3’ (Geiger et al., 2014) for COI and PCR primers for *Cytb* and *Act-2*. The obtained sequences were deposited into the GenBank database (see Table S1 for accession numbers).

### 2.2. Analysis of mtDNA data

The nucleotide sequences of *Cytb* and *CO*I were manually aligned, edited, and checked for unexpected stop-codons using SeaView 4.4.2 (Gouy et al., 2010). The sequences were collapsed into haplotypes. Phylogenetic reconstruction was performed using the maximum-likelihood (ML) and the Bayesian inference (BI) approaches on each marker, both separately and combined. Before running phylogenetic analyses, the datasets were tested for redundancy and saturation using METAPIGA 3.01 (Helaers and Milinkovitch, 2010). We determined the best fit models of nucleotide substitution and the optimal partitioning scheme for each marker and combined the dataset using either IQ-TREE 1.5.4 (Nguyen et al. 2015; Kalyaanamoorthy et al. 2017) or PartitionFinder 2.1.1 (Lanfear et al., 2012) under the Bayesian Information Criterion (BIC). The partition schemes selected by IQ-TREE (Table 2) were subsequently used in the ML search with the same software, using 1 000 ultrafast bootstrap replicates (Minh et al., 2013).

**Table 2.** The best partition schemes.

| Partitions | *Cytb* | *COI* | Concatenated *Cytb*+*COI* |  |
| --- | --- | --- | --- | --- |
| ML |  |  |  |  |
| 1 Pos. | TN+I+G | TIM+I+G | *Cytb*: TIM2+G | *COI*: HKY+I |
| 2 Pos. | TN+I+G | TIM+I+G | *Cytb*: HKY+I | *COI*: HKY+I |
| 3 Pos. | TN+I+G | TIM+I+G | *Cytb*: HKY+I | *COI*: TIM2+G |
| BI |  |  |  |  |
| 1 Pos. | K80+I+G | K80+I+G | *Cytb*: HKY | *COI*: K80+I+G |
| 2 Pos. | HKY+G | F81+I | *Cytb*: K80+I+G | *COI*: HKY |
| 3 Pos. | GTR+G | GTR+G | *Cytb*: GTR+G | *COI*: GTR+G |

The following settings were common for all MrBayes runs: the sampling frequency was every 500 generations, and the first 25 % of generations were discarded as burn-in. Convergence of runs was assessed by examination of the average standard deviation of split frequencies and the potential scale reduction factor. In addition, stationarity was confirmed by examining the posterior probability, log likelihood, and all model parameters by the effective sample sizes (ESSs) in the program Tracer v1.6 (Rambaut et al., 2014). The phylogenetic trees resulting in ML and BI analyses were visualized and edited using FigTree v.1.4.3 (Rambaut, 2008).

The number of haplotypes (H), haplotype diversity (h), nucleotide diversity (*π*), number of segregating sites (S), and average number of nucleotide substitutions (K) were calculated using DnaSP v.5.10 (Librado and Rozas, 2009). Haplotype networks were built with the help of PopART 1.7 software (Leigh and Bryant, 2015) using the median-joining algorithm (Bandelt et al., 1999).

Divergence times between the *Barbus* species were estimated in BEAST 1.8 (Drummond et al., 2012) using a relaxed molecular clock with an uncorrelated lognormal distribution and a Yule speciation prior (Drummond et al., 2006). *Cytb* sequences of the *Barbus* species set were used for phylogeny as well as for comparative sets of the *Capoeta* and *Luciobarbus* species, some sequences of which were obtained from GenBank (Table S2). A total of 215 sequences with the length of a fragment of *Cytb* = 993 bp were analyzed. Outgroups were *Cyprinus carpio* (AY347287), *Cyprinion macrostomus* (AF180826), and *Aulopyge huegelii* (AF287416). The following calibration points were selected: (1) fossil evidence of the *Barbus* species (18–19 Mya, Czech Republic, several localities - Böhme and Ilg, 2003); (2) fossil evidence of the *Luciobarbus* genus (16.9–17.7 Mya, Turkey, Belenyenice - Böhme and Ilg, 2003); (3) fossil evidence of the Iberian Peninsula *Luciobarbus* species (5.33–7.05 Mya) (Doadrio and Casado, 1989; García-Alix et al., 2008). The substitution models by codon position were selected in PartitionFinder 2.1.1: K80+I+G, HKY+I, TN93+G. The user-specified starting tree was used. The tree was generated with MrBayes and then transformed into an ultrametric, time-calibrated tree using a penalized likelihood approach (Sanderson, 2002) implemented using the ‘chronos’ function of the ‘ape’ R package (Paradis et al., 2004). Each final MCMC chain was run for 10^8^ generations (20 % burn-in), with parameters sampled every 1 000 steps. Tracer 1.6 (Rambaut and Drummond, 2013) was used to evaluate the run convergence and effective sample sizes for all parameters. The trees were summarized with TreeAnnotator 1.7 (Drummond et al., 2012) to obtain a maximum clade credibility tree.

### 2.3. Analysis of nDNA data

Heterozygotes of individual sequences of *Act-2* were identified by double peaks in the chromatograms. All polymorphic sites were carefully inspected to ensure consistent identification of such positions. The sequences were aligned using Muscle 3.8.31 (Edgar, 2004). Alleles (haplotypes) of the *Act-2* were inferred using Phase version 2.1 (Stephens and Donnelly, 2003; Stephens et al., 2001) with 1 000 burn-in steps and 1 000 iterations. Some individuals carried alleles of different sizes due to insertion/deletion polymorphisms, as shown by the offset chromatogram peaks. Due to this constraint of using software that is commonly used for phase data, haplotypes were thus manually phased using the method described by Sousa-Santos et al. (2005) and Flot et al. (2006). The phylogenetic network between studied species was resolved by a statistical parsimony algorithm implemented in TCS 1.21 (Clement et al., 2010). The method was selected due to its ability to handle indels (gaps) as 5^th^ state. However, the software considers each gap position within an alignment as a single mutation, which is a very implausible scenario for the origin of indels. To overcome this shortcoming, we cut all deletions within the alignment to the length of one bp. After such modification, each indel was handled by TCS as a single mutation event, which creates a much more probable model. Given that clipped fragments did not contain any nucleotide substitutions, this approach allowed us to use all sources of variability without any information loss. The nucleotide variability of *Act-2* sequences were described in the same way as for the mtDNA loci. Both alleles for each individual were accounted for in all analyses.

## 3. Results

### 3.1. Mitochondrial DNA sequence analyses

Based on the *Cytb* sequences, 27 studied species of the *Barbus* s. str., including the Caucasian taxa from the monophyletic cluster that is sister to the *Aulopyge, Luciobarbus* and *Capoeta* species group (Fig. 2), confirmed the results of previous phylogenetic studies (Levin et al., 2012; Machordom and Doadrio, 2001; Tsigenopoulos et al., 2003). One individual of the Sevan barbel *B. goktschaicus* had a haplotype of the algae eater, Sevan khramulya *C. capoeta* (Fig. 2, Fig. S1), which is an evolutionary hexaploid (2n=150, see Krysanov, 1999), suggesting introgressive hybridization between the tetraploid *Barbus* and hexaploid *Capoeta* lineages.

**Fig. 2.**
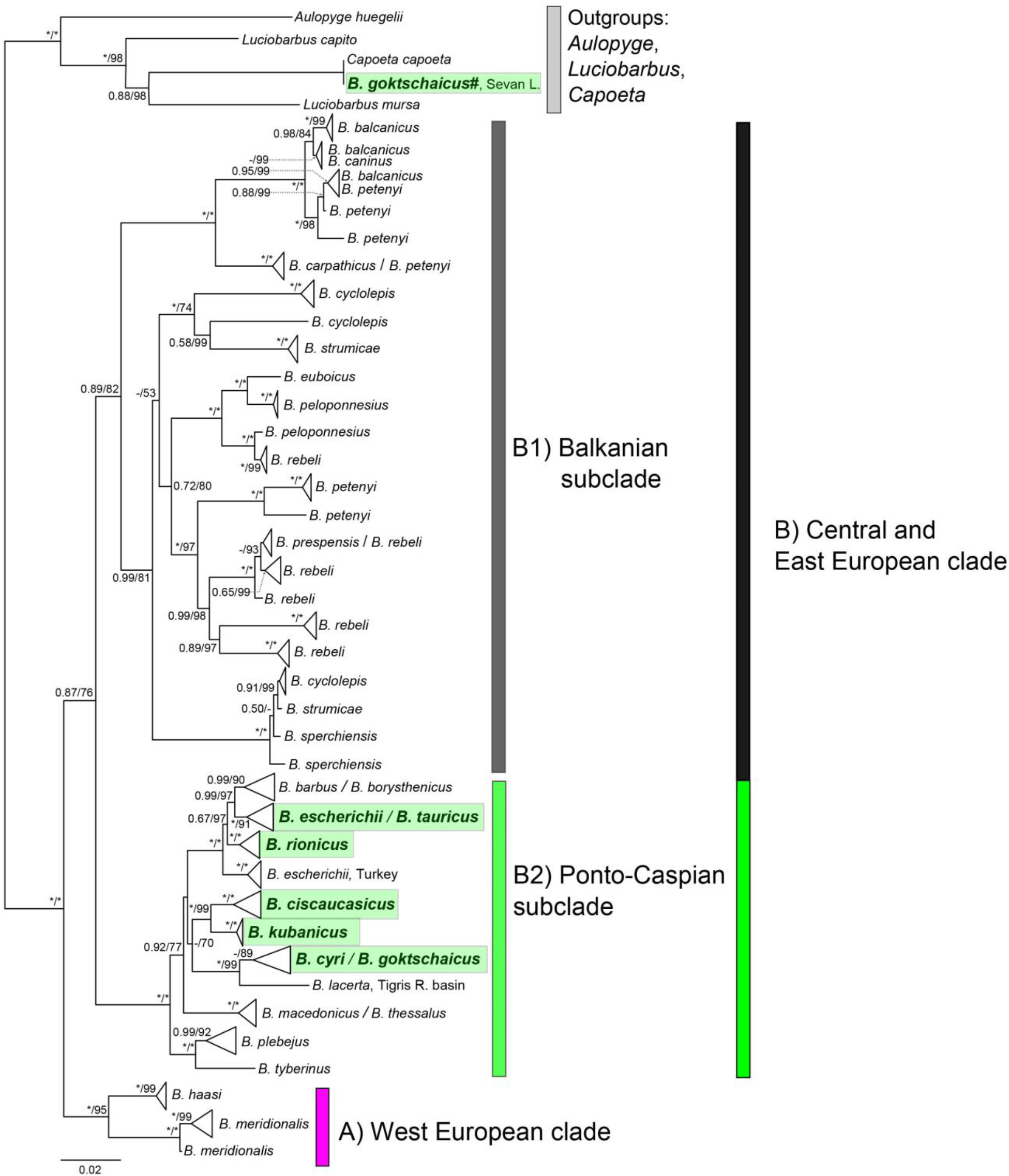
BI consensus gene tree based on the partial *Cytb* sequences (993 bp long) for 27 *Barbus* species. Bayesian posterior probabilities (on the left side) and bootstrap values from a ML analysis (on the right side) above 0.5/50 are shown; asterisks represent posterior probabilities/bootstrap values of 1/100. Caucasus taxa are highlighted in green. The nodes with multiple specimens were collapsed to a triangle, with the horizontal depth indicating the level of divergence within the node. One individual of *B. goktschaicus* had the same haplotype as *C. capoeta* and considered as a hybrid (marked with #). Scale bar and branch lengths are in expected substitutions per site.

According to both ML and BI analyses, the genus *Barbus* is subdivided into two main clades: A) the West-European clade with two species - *B. haasi* and *B. meridionalis* and B) the remaining species from Central and Eastern Europe. This last group (B) is composed of two subclades: B1) the largest and most diversified Balkanian group of barbels, with moderate support, which includes 12-13 species, mostly from the Balkan peninsula, and *B. caninus* from Northern Italy; and B2) the Ponto-Caspian subclade that includes all Caucasian species, one species from the Tigris-Euphrates basin (*B. lacerta*), two species from the Balkan Peninsula (*B. macedonicus* and *B. thessalus*) and two species (*B. plebejus* and *B. tyberinus*) distributed in the Apennine Peninsula and the Western part of the Balkan Peninsula. Within the last subclade, two main monophyletic groups were recovered, one with Ponto-Caspian species and the other with the species of the Italian and Western Balkanian area. The most basal group among all *Barbus* spp. is composed of the mainly Iberian species, *B. meridionalis* and *B. haasi*.

Data on the *CO*I marker confirmed the presence of the main inner clades of *Barbus* that was revealed by the *Cytb* analyses; however, the phylogenetic utility of the *CO*I marker to resolve inner relationships was low (Table S3, Fig S1). The combined dataset *Cytb+COI* analyses confirmed the presence of main clades in *Barbus* and its sister relationships to *Luciobarbus* + *Capoeta*, which were revealed by the *Cytb* marker.

### 3.1.1. Intrarelationships of Ponto-Caspian subclade

The Ponto-Caspian subclade includes 11 lineages distributed mainly in the basins of the Black Sea [5 lineages: i) *B. barbus* + *B. borysthenicus;* ii) *B. tauricus* + *B. escherichii;* iii) *B. rionicus;* iv) *Barbus* sp. from Turkey, and v) *B. kubanicus* – all these lineages were numbered according to the lineages delineated by Kotlík et al. (2004) for Black Sea drainage barbels] and the Caspian Sea [3 lineages: vi) *B. ciscaucasicus*, and vii) *B. cyri* + *B. goktschaicus*, as well as viii) *B. lacerta* lineage from the drainage of the Persian Gulf, Arabian Sea basin], ix) the Aegean lineage that comprises *B. macedonicus* and *B. thessalus* from the Aegean Sea drainage, x) the *B. plebejus* lineage distributed in the Western Balkans and in the Apennine Peninsula, and xi) the *B. tyberinus* lineage from the Apennine Peninsula (Fig. 2).

**Fig. 3.**
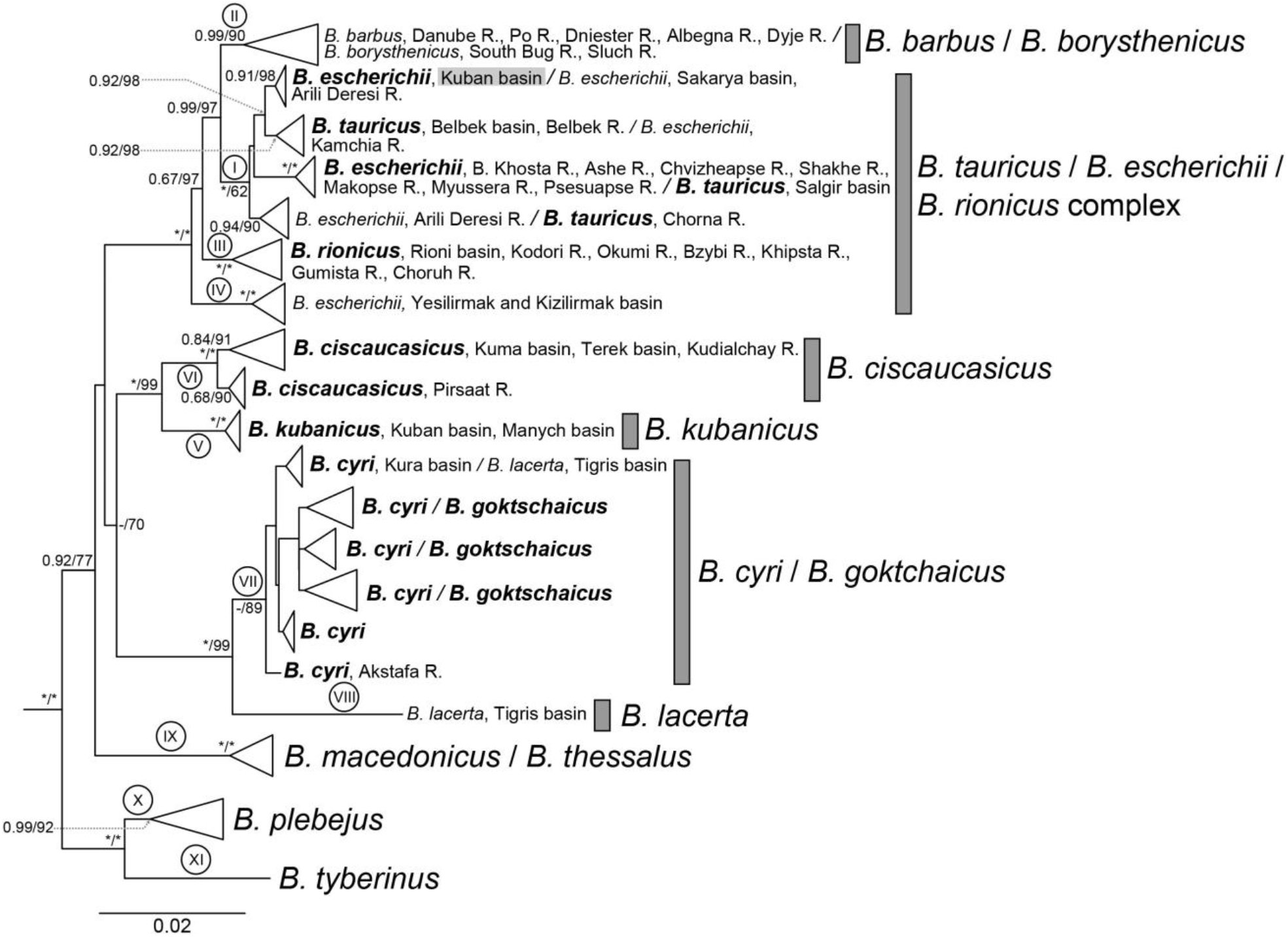
The enlarged fragment of BI consensus gene tree (Fig. 2) displayed the Ponto-Caspian species inside B2 subclade based on the 69 unique haplotypes of *Cytb* sequences (993 bp long). Bayesian posterior probabilities (on the left side) and bootstrap values from a ML analysis (on the right side) above 0.5/50 are shown; asterisks represent posterior probabilities/bootstrap values of 1/100. Lineages from I to V corresponds to such in study of Kotlík et al. (2004). The species *B. escherichii* detected in the Kuban basin is highlighted with grey. Scale bar and branch lengths are in expected substitutions per site.

The Caucasian barbel species are clustered into three groups: a) the Black Sea group, which includes all Black Sea species, except for *B. kubanicus*; b) the Caspian group, which includes *B. cyri* with *B. goktschaicus* (the latter does not differ from the former genetically - see Table S3; both inhabit the Kura-Araks system), and *B. lacerta*, which is the sister to *B. cyri* and inhabits the Tigris-Euphrates drainage adjacent to the Kura-Araks drainage, and c) the Ciscaucasian group, which is composed of *B. ciscaucasicus*, which inhabits some northern tributaries of the Caspian Sea, and *B. kubanicus*, which inhabits the Kuban basin (Azov Sea drainage) and is sister to *B. ciscaucasicus*. All of the Caucasian species combined are sister to the group of the two Aegean species: *B. macedonicus* and *B. thessalus*. The Aegean lineage is deeper diverged compared to the Caucasian lineages. The outer group in the Ponto-Caspian subclade is *B. plebejus+B. tyberinus*, which is a westward distributed species.

The divergence between members of the Ponto-Caspian subclade is less than between species of another large subclade, the Balkanian subclade (Fig. 2), which suggests the younger origin of the Ponto-Caspian species (see section 3.3. Molecular clock).

### 3.1.2 Phylogeography of Black Sea coastal B. tauricus / escherichii / rionicus complex

Populations of the Black Sea group have a rather complicated phylogeography. This group is characterized by 24 *Cytb* haplotypes subdivided into three main haplogroups (Fig. 4). The most diverged haplogroup includes the *B. tauricus* / *B. escherichii* complex with 12 haplotypes. The Crimean barbel *B. tauricus*, with a restricted range to rivers in the Crimean Peninsula, has only one shared haplotype with barbels from other populations. Despite that, its haplotypes are very divergent among each other and are closer to haplotypes from the geographically remote populations of *B. escherichii* (Black Sea tributaries in Turkey and Bulgaria) or closely located ones (rivers of Russian coast of Black Sea) (Fig. 5). The species *B. tauricus* also had the highest nucleotide diversity (Table 3).

**Fig. 4.**
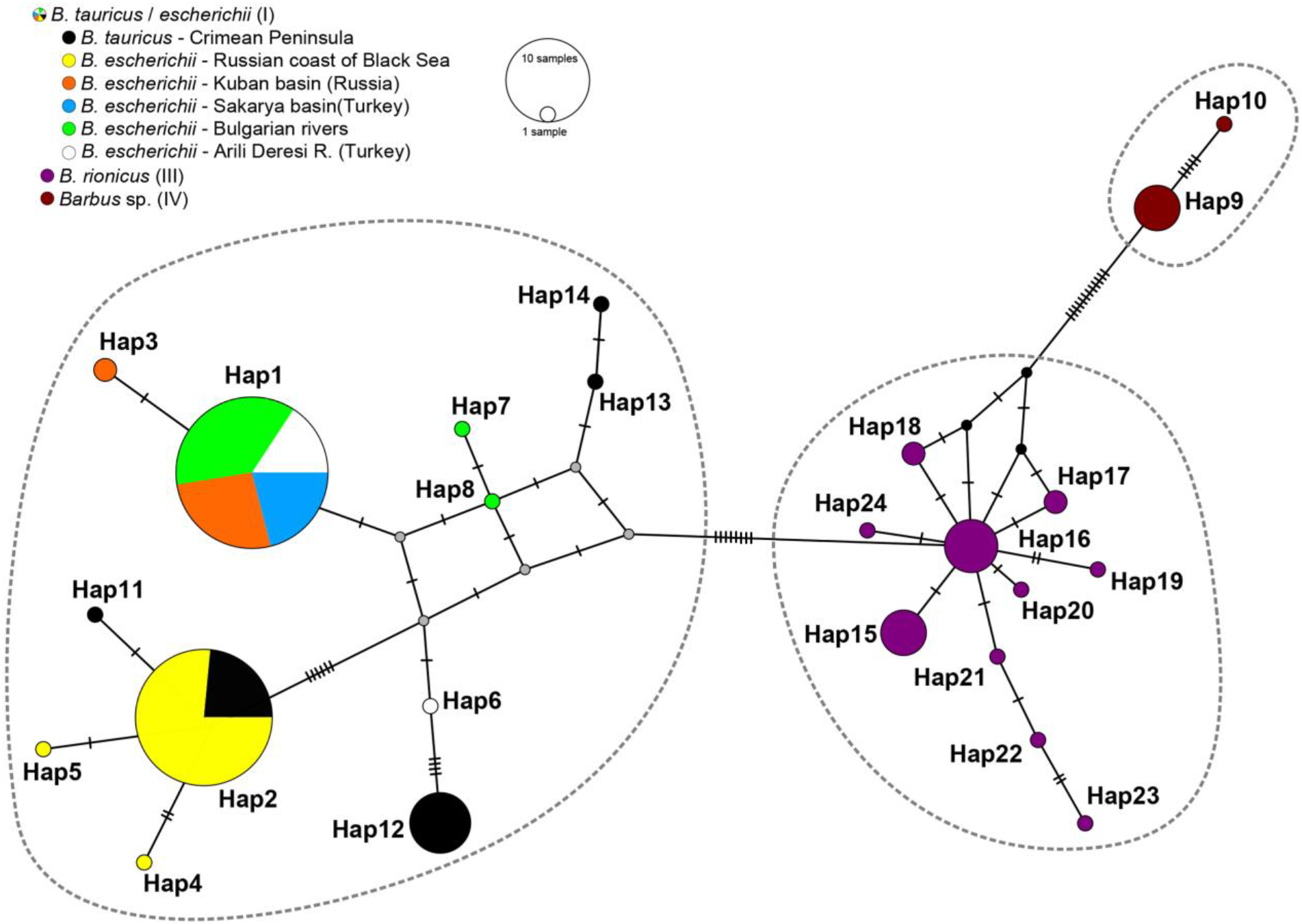
Median-joining haplotype network of Black Sea barbs *B. tauricus* / *escherichii* / *rionicus* complex constructed on *Cytb* sequences. The circle sizes correspond to relative haplotype frequencies. One vertical mark denotes one substitution. Small grey circles represent hypothetical intermediate haplotypes.

**Fig. 5.**
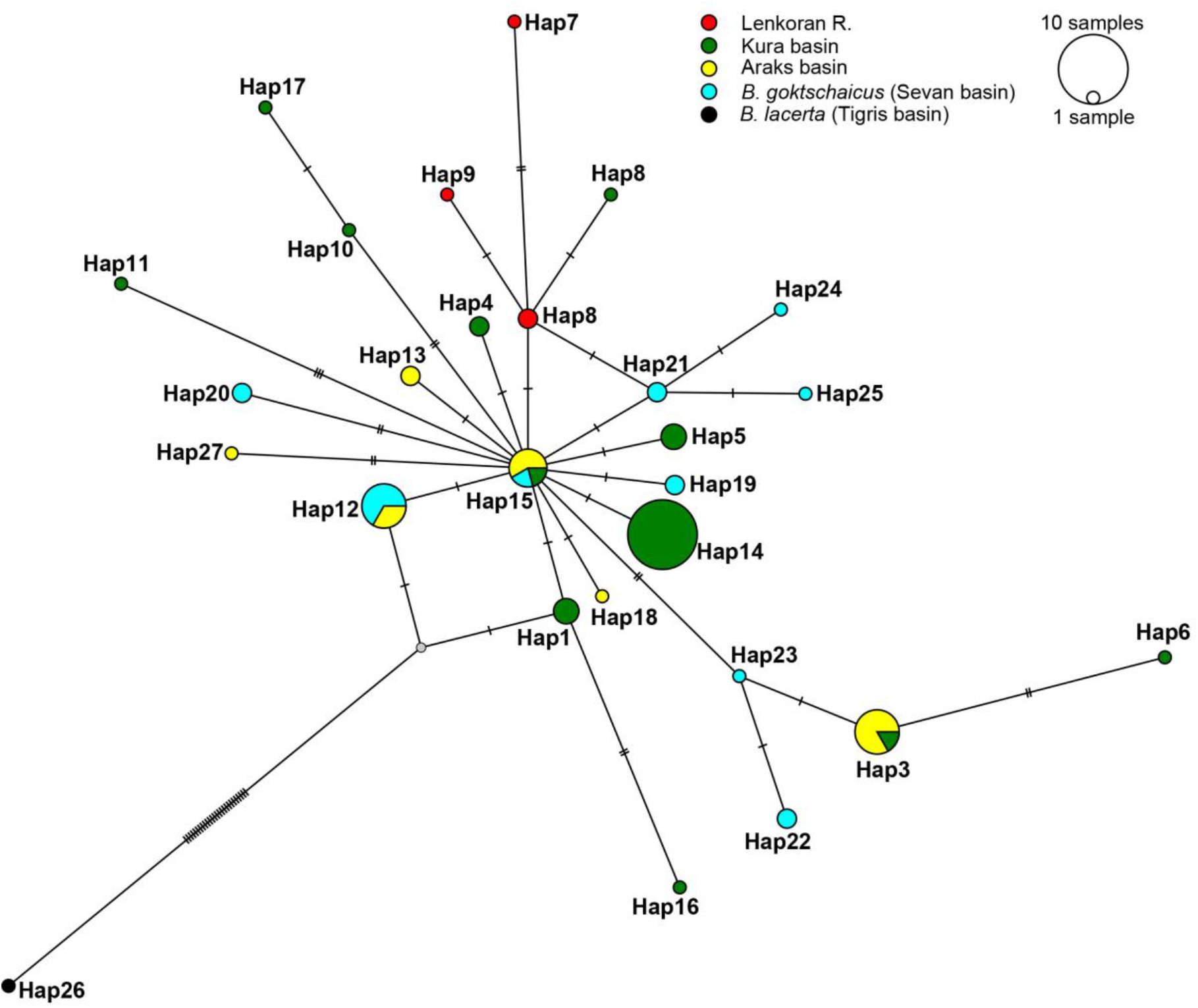
Median-joining haplotype network of *B. cyri* / *goktschaicus* constructed on *Cytb* sequences. The circle sizes correspond to relative haplotype frequencies. One vertical mark denotes one substitution. Small grey circles represent hypothetical intermediate haplotypes.

**Table 3.**
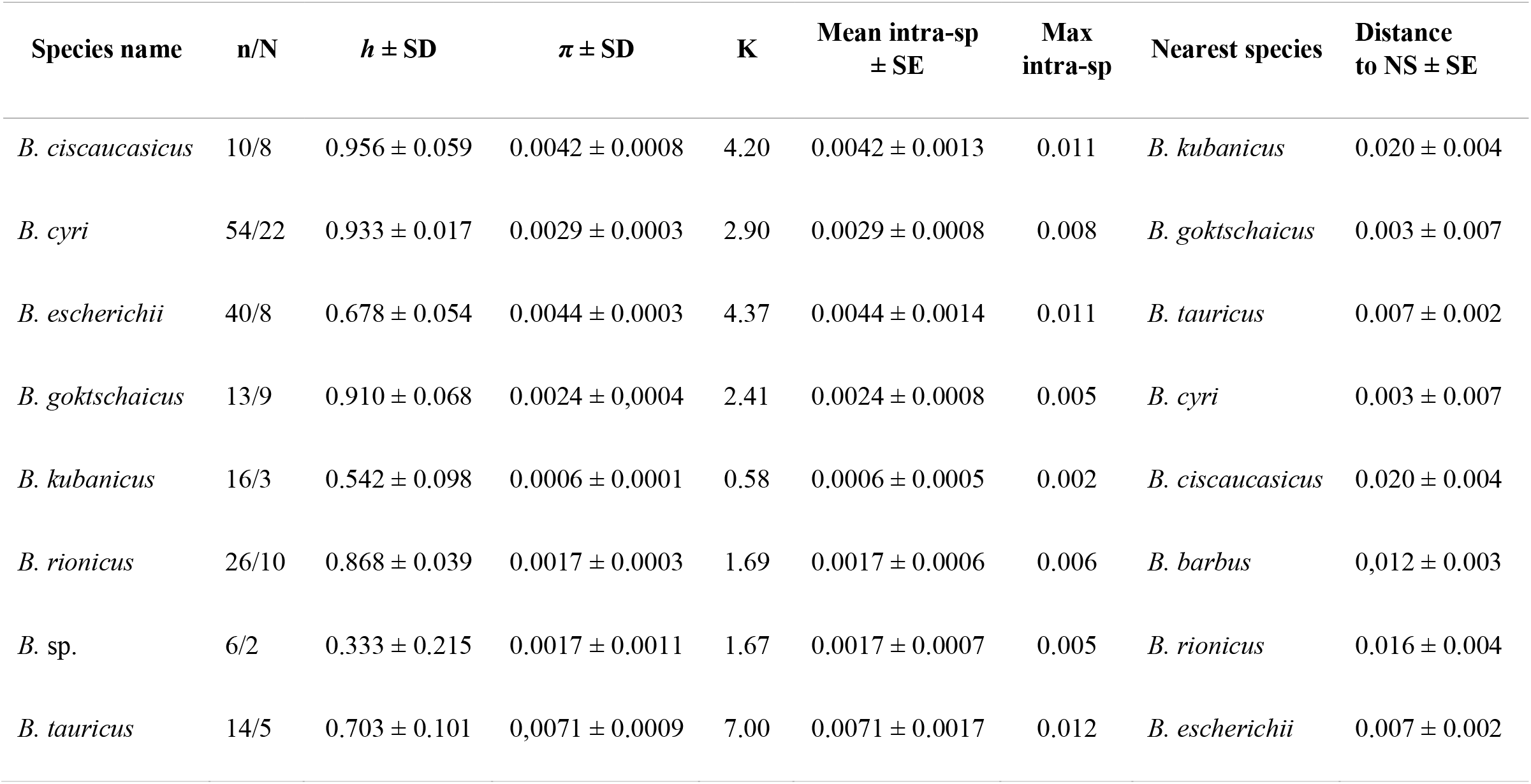
Intraspecific genetic variation of *Barbus* spp. from Caucasus region and Crimean Peninsula in *Cytb* sequences. n/N - sample size/number of haplotypes; *h* ± SD - haplotype diversity ± standard deviation; *π* ± SD - nucleotide diversity (per site) ± standard deviation; K - average number of nucleotide differences; Mean intra-sp ± SE - mean intraspecies p-distance ± standard error; Max intra-sp - maximum intraspecies p-distance; Distance to NS ± SE - mean p-distance to a nearest species ± standard error (based on 1000 bootstrap replications).

We have to note that *B. escherichii* has been detected in the Kuban basin for the first time, where from only *B. kubanicus* was previously reported. Remarkably, the Kuban haplotypes of *B. escherichii* are closer to or share the same haplotypes with populations from the southern coast of the Black Sea in Turkey (the Sakarya basin and the Arili Dereseli R. near Rize city) but are adjacent to the *B. escherichii* from the Russian coast of Black Sea (the rivers between Sochi and Tuapse).

There are two other distinct haplogroups, which are distant to *B. tauricus/escherichii* complex and to each other, and which represent *B. rionicus* and *Barbus* sp. (assigned as lineage iv in study of Kotlík et al., 2004). The species *B. rionicus* is widely distributed from the Choruh basin along the Georgian coast to the Bzyb River and has a star-like pattern of 10 haplotypes (Fig. 1 & 5). The most diverged haplogroup represents two haplotypes from the center of the Southern coast of the Black Sea in Turkey (Kizilirmak and Yesilirmak basins) and are assigned here as *Barbus* sp.

### 3.1.3. Phylogeography of the Kura barbel Barbus lacerta and Sevan barbel B. goktscaicus

The Sevan barbel *B. goktschaicus* is represented by nine haplotypes (h=0.910), two of which are shared with haplotypes of *B. cyri* (Fig. 5). The other seven haplotypes are not divergent from *B. cyri*, which is represented by 21 haplotypes (h=0.933). The P-distance between these two species is minute (0.003). The most divergent haplotype of *B. cyri* (8 mutational steps from the ancestral haplotype) was detected from the Qarachai R. (loc. 30), a northern tributary of the Kura R. The species *B. lacerta* inhabited the Tigris-Euphrates system and, being sister to *B. cyri*, had 28 mutational steps to the nearest haplotype of the latter.

### 3.2. Intron 2 beta actin data set

The statistical parsimony network generated by TCS is not very diversified (Fig. 6). All Caucasian barbel species are characterized by 16 unique alleles of *Act-2*. The most frequent allele has an ancestral position and was formed by individuals of the Caucasian species *(B. cyri, B. goktschaicus* and *B. escherichii)* and some individuals of the Balkanian species (white color). Moreover, this is the only allele shared between the Balkanian and Ponto-Caspian subclades. More diversified representatives of the Balkanian subclade, with 26 different alleles of six species (sample size 40 individuals) compared to 16 alleles of six Caucasian and one Crimean species (52 individuals) coupled with mtDNA data, suggest a younger origin and shallower intra-divergence of the Ponto-Caspian subclade.

**Fig. 6.**
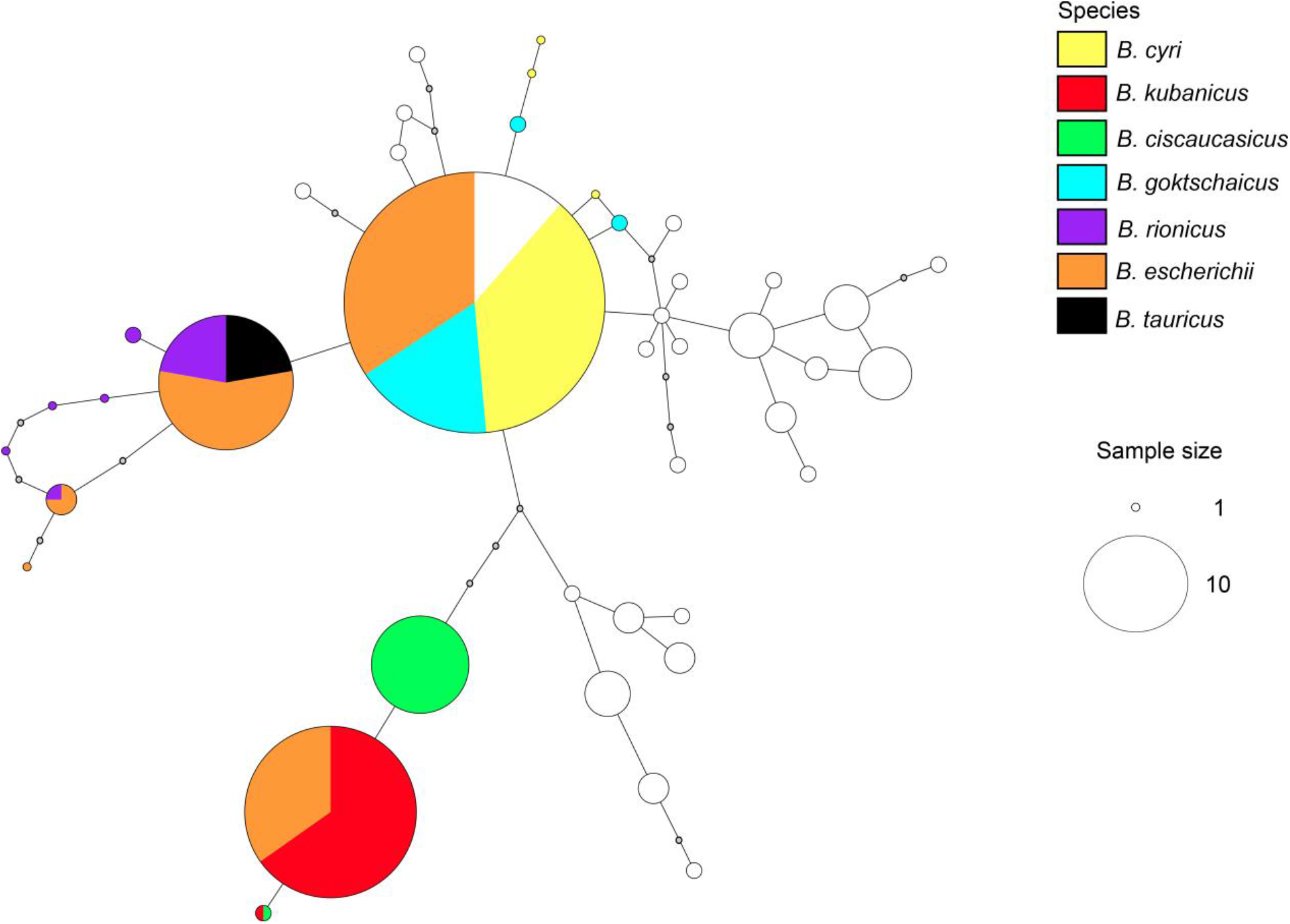
Statistical parsimony network of Caucasian barbels constructed on *Act-2* phased sequences. Empty circles represent alleles of Balkanian species *B. balcanicus*, *B. petenyi*, *B. prespensis*, and *B. rebeli* from study of Antal et al. (2016). The circle sizes correspond to relative allele frequencies. Small grey circles represent intermediate mutational steps.

**Table 4.**
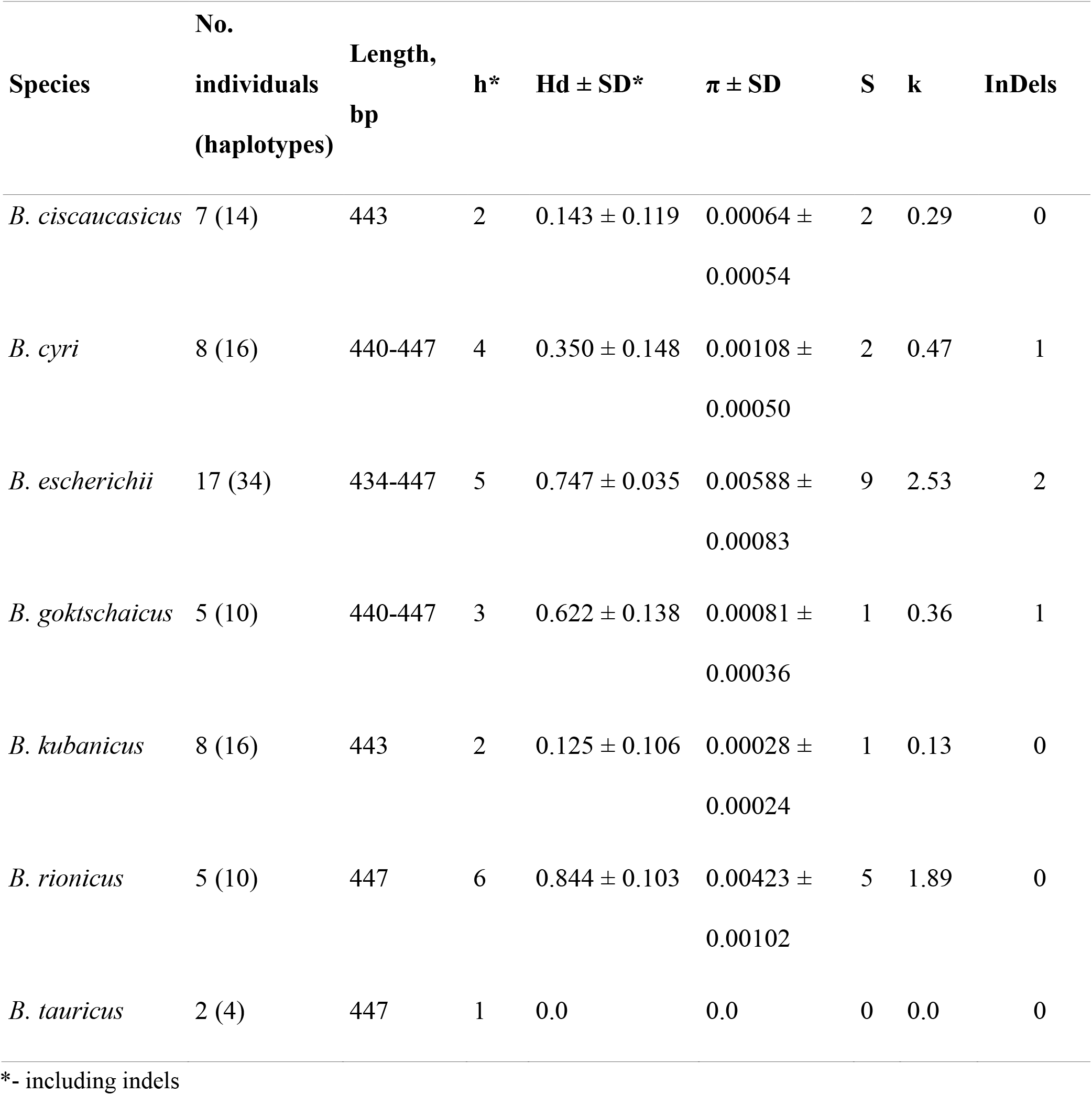
Intraspecific genetic variation of Caucasian barbels in nuclear marker Act-2.

| Species | No.<br>individuals<br>(haplotypes) | Length,<br>bp | h* | Hd ± SD* | $\pi \pm$ SD | S | k | InDels |
| --- | --- | --- | --- | --- | --- | --- | --- | --- |
| *B. ciscaucasicus* | 7 (14) | 443 | 2 | 0.143 ± 0.119 | 0.00064 ±<br>0.00054 | 2 | 0.29 | 0 |
| *B. cyri* | 8 (16) | 440-447 | 4 | 0.350 ± 0.148 | 0.00108 ±<br>0.00050 | 2 | 0.47 | 1 |
| *B. escherichii* | 17 (34) | 434-447 | 5 | 0.747 ± 0.035 | 0.00588 ±<br>0.00083 | 9 | 2.53 | 2 |
| *B. goktschaicus* | 5 (10) | 440-447 | 3 | 0.622 ± 0.138 | 0.00081 ±<br>0.00036 | 1 | 0.36 | 1 |
| *B. kubanicus* | 8 (16) | 443 | 2 | 0.125 ± 0.106 | 0.00028 ±<br>0.00024 | 1 | 0.13 | 0 |
| *B. rionicus* | 5 (10) | 447 | 6 | 0.844 ± 0.103 | 0.00423 ±<br>0.00102 | 5 | 1.89 | 0 |
| *B. tauricus* | 2 (4) | 447 | 1 | 0.0 | 0.0 | 0 | 0.0 | 0 |

\*- including indels

The Ciscaucasian species *B. ciscaucasicus* and *B. kubanicus* are distinct from other species (four mutational steps to both ‘ancestral’ alleles or Balkanian species). These species are close to each other and are rather monomorphic (Table 4), having two main species-specific alleles and one allele shared by a single individual of both species (Fig. 6). The shared haplotype could suggest both hybridization in the past or a retention of ancestral polymorphism due to incomplete lineage sorting (ILS).

The sharing of the same allele by *B. kubanicus* and *B. escherichii* recorded in the Kuban basin provides evidence for the hybridization between these species rather than ILS, since other populations of the *B. escherichii* / *tauricus* / *rionicus* complex had no shared alleles with *B. kubanicus* (Fig. 6). Notably, all studied individuals of *B. escherichii* that penetrated into the Kuban basin had species-specific mtDNA haplotypes (Fig. 4), while six of them were heterozygous in *Act-2* and possessed one allele specific to *B. kubanicus*, an endemic species of the Kuban river system (Fig. 6). Furthermore, four individuals in the Kuban population were assigned to *B. escherichii* based on their mtDNA being homozygous with the allele specific to *B. kubanicus*, i.e., were fully introgressed.

The complex *B. tauricus* / *escherichii* / *rionicus* is the most diversified by the nuclear marker representing eight alleles, excluding the supposed hybrids from the Kuban R. This diversification, however, provides no evidence of the genetic subdivision of the species of this complex based on *Act-2* and suggests that the recent origin and likely secondary contacts were between isolated and semi-isolated populations.

### 3.3 Molecular clock

Divergences between the well-established genera *Barbus* and *Luciobarbus* were estimated to have occurred in the Late Oligocene-Early Miocene sub-epoch (95 % HPD: 19.26-24.43 Mya – Fig. 8), which is close to the previously obtained results (Ghanavi et al., 2016; Levin et al., 2012). A separation of the Ponto-Caspian subclade from the Balkanian subclade was estimated to occur during the Middle Miocene sub-epoch approximately 16.94 Mya (95 *%* HPD: 14.82-18.76 Mya). The divergence of the Balkanian species of the Ponto-Caspian subclade *(B. macedonicus* and *B. thessalus* lineage) and of the Ponto-Caspian species themselves was estimated to have occurred in the Late Miocene sub-epoch approximately 10.26 Mya (95 % HPD: 7.12-13.58 Mya). The lineage of Ponto-Caspian barbels was early diverged for Black Sea and Caspian Sea groups approximately 9.45 Mya (95 % HPD: 6.46-12.50).

## 4. Discussion

### 4.1. Phylogeny of the genus Barbus and position of Caucasian barbels

The phylogenetic relationships of the genus *Barbus* s. str. have been reported in several studies based on the *Cytb* marker (Berrebi and Tsigenopoulos, 2003; Kotlík and Berrebi, 2002; Zardoya and Doadrio, 1999). However, all Caucasian barbels represented by seven species were unstudied. The phylogeny constructed by BI and ML approaches in the present study is based on the mtDNA sequences of 27 species of *Barbus* s. str., which are clustered for two clades. The earlier diverged clade is the West European clade with two species, *B. meridionalis* and *B. haasi*. The presence of this clade was shown in previous phylogenetic studies, however, without separation and naming (Berrebi and Tsigenopoulos, 2003; Zardoya and Doadrio, 1999). The second, much diversified clade of barbels from Central and East Europe includes all other species with deeply diverged lineages, especially in the Balkans. The inner phylogeny of the Balkanian subclade deserves special attention and should be studied more thoroughly, which is beyond the scope of the present study.

The diversified Balkanian subclade is sister to the Ponto-Caspian subclade. The divergence of these subclades probably occurred ca. 17 Mya, according to our molecular clock (Fig. 7), during one of the Alpine orogeny phases and was accommodated by a continuing collision of the Afro-Arabian plate with the Eurasian plate (Jolivet and Facenna, 2000). This date is close to that which was reported in a previous study (15.7 Mya), which was based on the restricted number of *Barbus* species (Levin et al., 2012). However, the divergence between Caucasian and West Balkan cave shrimp *Troglocaris* was estimated to occur earlier, at approximately 6-11 Mya based on COI and 16S markers (Zakšek et al., 2007). Earlier diverged lineages inside the Ponto-Caspian subclade are represented by lineages of *B. tyberinus* and *B. plebejus*, which are distributed in the Apennine Peninsula and South-Western Balkans. They are sister to nine other lineages, seven of which are Caucasian, that are distributed in the basins of the Black and Caspian Seas. Two other lineages branched earlier, including the Aegean species *(B. macedonicus* and *B. thessalus* – lineage ix) and the Mesopotamian species (*B. lacerta* from the Tigris-Euphrates basin Arabian Sea drainage – lineage viii).

**Fig. 7.**
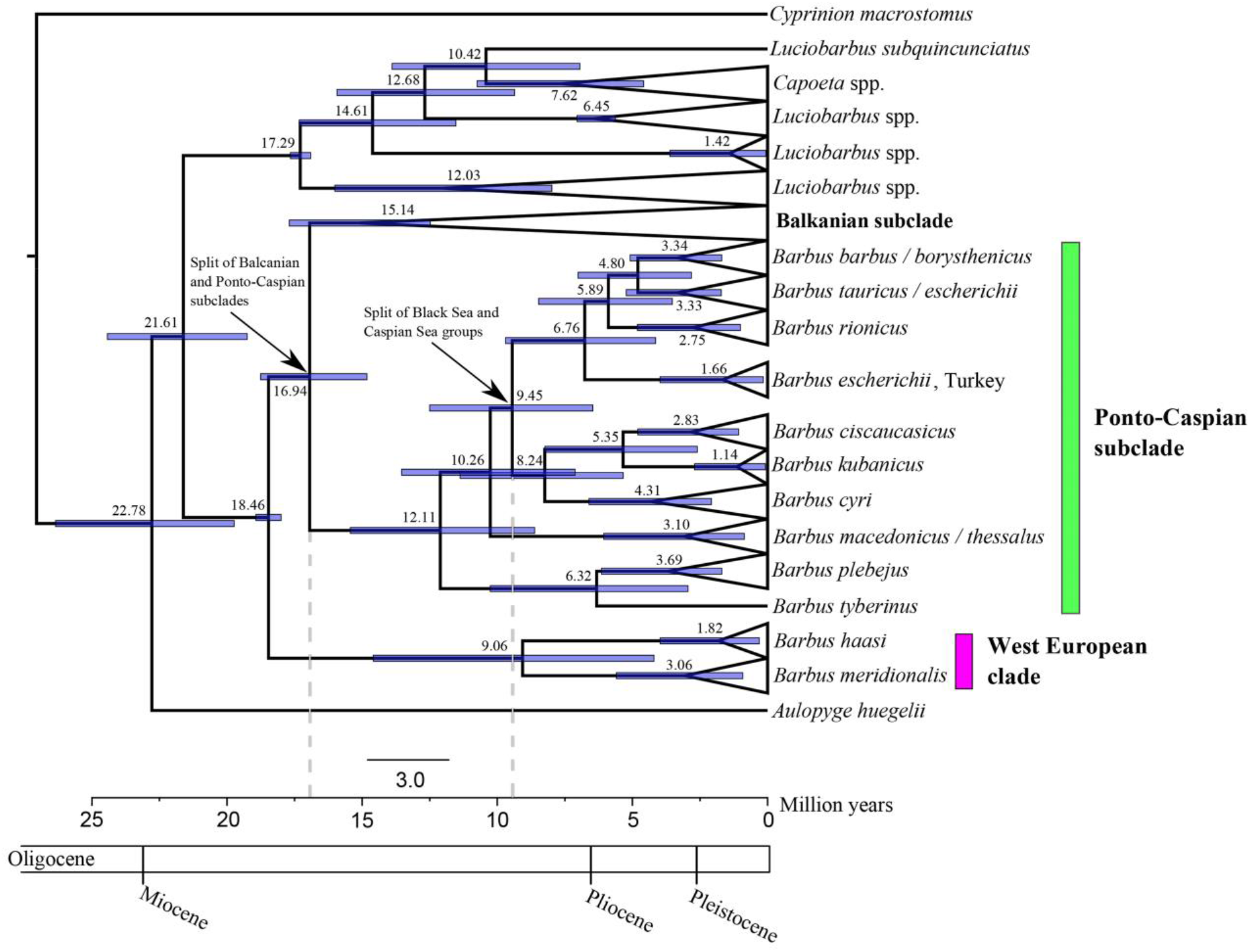
Divergence time (Mya) and credibility intervals (95% highest posterior density). All nodes were highly supported at PP=1 (not shown on tree).

The present study confirmed the relatedness of two Balkanian barbel species that inhabited the Aegean Sea drainage to Caucasian barbels (Levin et al., 2012) in the Ponto-Caspian subclade. The Balkanian drainage of the Aegean Sea shares a watershed with the western part of the Black Sea drainage. There are genetic evidence of the close relationships between Caspian lamprey *Caspiomyzon wagneri*, which inhabited the Caspian Sea and the ‘*Eudontomyzon’ hellenicus*, which inhabited the Struma basin (Lang et al., 2009), or even the occurrence of Ponto-Caspian fish species in the water bodies of the Aegean Sea drainage (Levin et al., 2017; Tsoumani et al., 2014).

Caucasian barbels are composed of two main lineages, Black Sea and Caspian lineages, which are represented mainly by species of the Black Sea drainage (except for *B. kubanicus)* and species of the Caspian Sea drainage (including *B. lacerta* from the Tigris-Euphrates) (Fig. 2). Based on early branching of these two main lineages, they were split early after the colonization of the Ponto-Caspian basin. According to our molecular clock, a separation of groups of Caucasian barbels of the Black and Caspian Seas groups occurred during the Lower Miocene sub-epoch approximately 9.45 Mya (Fig. 7), which was a period of intensive elevation of the Caucasian region and occurred when the modern relief was forming (Lazukov, 1989). The Suram (or Likhi) range in Transcaucasia that separated the drainages of the Black and Caspian seas raised ca. 10-15 Mya and has fully isolated the Western and Eastern parts of Transcaucasia ca. 10 Mya (Maruashvili, 1981). The Caspian lineage is more diversified and includes three groups: a) the Mesopotamian species *B. lacerta*, b) the Eastern Transcaucasian species *B. cyri* / *goktschaicus*, and c) the Ciscaucasian sister species *B. ciscaucasicus* and *B. kubanicus*. The intra-Caucasian divergence occurred from 6.76 to 4.80 Mya based on our molecular clock (Fig. 7), which coincides with a period of active uplift of both the Great and Lesser Caucasus, as well as the Pliocene volcanism in Transcaucasia (Lazukov, 1989). The possible reasons for shallower divergence between the Circum Black Sea species will be considered below.

Based on the phylogeny of the *Barbus* s. str. and the degree of diversification among the different subclades, we hypothesize that the genus could have originated from the common ancestor of *Luciobarbus*/*Barbus*, whether in A) Western Europe and further dispersed into Central and Southern Europe (mainly Balkans), or in B) Balkanian Europe and further distributed throughout Europe and penetrated into Minor Asia and Caucasus. The Balkan region is a European biodiversity hotspot that was suggested as a refuge area for fish species (Oikonomou et al., 2014; Tsigenopoulos, 1999; Zardoya et al., 1999). The West European clade branched earlier compared to other clades; however, i) differences in the branching distance are not large, ii) the Balkanian clade is much more diversified, and iii) the most ancient Balkanian species could become extinct during some of climatic oscillations or geological events. The latter reason indicates that the West European species might look more ancient. Currently, we cannot provide support any of these hypotheses on the origination of the genus until we obtain more data.

### 4.2. Hybridization and introgression

Interploidy hybridization between hexaploid *C. capoeta sevangi* (2n=150 - Krysanov, 1999) and tetraploid *B. goktschaicus* (2n=100 - Krysanov, 1999) in Lake Sevan (Armenia) was recognized in the phenotypically *B. goktschaicus* individual that possessed the mtDNA of *C. capoeta sevangi*. Moreover, the short fragment of the nuclear marker *Act-2* of *Capoeta* was also detected in this hybrid specimen. It is the first evidence of the hybridization between these genera. Natural intergeneric hybrids between *Capoeta damascina* (2n = 150 according to Gorshkova et al., 2002) and *Luciobarbus longiceps* (not karyotyped, but should be a tetraploid based on its phylogenetic position – see Tsigenopoulos et al., 2003) were repeatedly reported from Lake Kinneret, Israel (Abraham, 1996; Stoumboudi et al., 1992, 1993). Interploidy hybridization is an apparent evolutionary mechanism for the origination of new polyploid fish lineages, including *Capoeta* (Chenuil, 1999; Levin et al., 2012; Yang et al., 2015), and the cases of such hybridization have to be studied more thoroughly.

While intergeneric hybrids are rarely recorded, interspecies hybridization inside the genus *Barbus* s. str. is a more common phenomenon. For instance, hybridization and introgression were detected between the species of the Western European clade *B. meridionalis* and *B. haasi* (Machordom et al., 1990), between *B. barbus* and *B. meridionalis* in France (Berrebi et al., 1993; Crespin et al., 1999), *B. barbus* and *B. carpathicus* in Slovakia (Lajbner et al., 2009), *B. barbus* and *B. plebejus, B. balcanicus* and *B. plebejus*, and *B. caninus* and *B. plebejus* in Italy (Buonerba et al., 2015; Meraner et al., 2013; Tsigenopoulos et al., 2002). In some cases, the asymmetrical hybridization with disproportionate contribution of *B. barbus* mothers was detected (Lajbner et al., 2009; Meraner et al., 2013). Moreover, in the case of *B. barbus* and *B. carpathicus*, no F2 progeny or backcrosses to *B. carpathicus* were found (Lajbner et al., 2009).

Hybridization between Ponto-Caspian barbels is reported for the first time in the present study. The case of hybridization between *B. escherichii* and *B. kubanicus* was established after finding *B. escherichii* in the Kuban basin (reported in this study for the first time). We interpret this event as a natural hybridization during secondary contact. The Kuban population of *B. escherichii* had mtDNA haplotypes similar to the remote conspecific Turkish populations. In our opinion, this is a relict population that resulted from one of the migration waves that occurred during the freshwater phases of the Black Sea (see a more detailed discussion in the subsection below). Remarkably, all sampled Kuban specimens of *B. escherichii* have their own species-specific mtDNA haplotypes, but a nuclear marker revealed that several individuals had the nuclear genome of *B. kubanicus*, and some were heterozygous. First, it gives evidence that hybridization occurred in the past and is ongoing (heterozygous specimens may be F1 hybrids or backcrosses). Second, it may reveal asymmetrical hybridization, as reported previously for several European species of barbels (Buonerba et al., 2015; Crespin et al., 1999; Lajbner et al., 2009; Meraner et al., 2013). The obtained results show that the hybridization of two barbel species in the Kuban basin needs further investigation.

### 4.3. Mosaic dispersal of lineages in Black Sea drainage

The spatial heterogeneity of the haplotype structure of different lineages in the Black Sea drainage is common for all studied taxa, except for *B. rionicus*, which inhabited the southeastern part of Black Sea basin and has unique haplotypes (Fig. 4, 6). In our view, the mosaic dispersal of mtDNA haplotypes is rather a consequence of the secondary contacts of different lineages during the freshwater phases of Black Sea over a span of the last 2 million years (Ryan et al.,2003). One such obvious example of colonization waves is the significantly diverged haplotypes in the rivers of the Crimean Peninsula. Crimean haplotypes were detected circum Black Sea in Bulgarian and Turkish rivers as well as in rivers of Russian coast of the Black Sea, from Sochi to Tuapse. The possibly mixed haplotypes in Crimea reflect the complicated history of the Crimean rivers, which changed their directions and mouth position significantly. For example, the longest Crimean river, the Salgir, is a type locality for *B. tauricus* and has several ancient channels (Sludskiy, 1953). The most distant mouths of these channels are separated by >800 km along the modern coastline. It might be important for the settlement of different migrating groups brought by different lacustrine currents. We have no data on the direction of the currents in ancient freshwater Black Sea. Nevertheless, the Crimean Peninsula, being largely beaked into the Black Sea and having a prolonged coastal line, could serve as a significant ‘trap’ for collecting barbels that migrate from both western or eastern parts of the giant freshwater lake.

Another example of several waves of migrations is the presence of two different haplogroups of *B. escherichii* in the northeastern part of the Black Sea basin. The *B. escherichii* individuals detected in the Kuban basin had the same haplotypes as conspecific population from Turkey near Rize (distance is ca. 620 km along the coastline) and shared no haplotypes with closely located populations from the Black Sea rivers of the Russian Coast (ca. 120 km distance). This Kuban population of *B. escherichii* may be considered a relict population, as a remnant of the previous wave of colonization in the riverine system where the endemic barbel *B. kubanicus* strongly predominates.

Here, we agree with suggestions that such a phylogeographical pattern most likely reflects the allopatric divergence of multiple ancestral populations (Kotlík et al., 2004) with subsequent contacts and, supposedly, hybridization and gene flow. In addition, the Azov-Black Sea is considered to be one of important glacial refugia for many freshwater fish species, such as the chub *Squalius cephalus* (Durand et al., 1999), the perch *Perca fluviatilis* (Nesbø et al., 1999), the brown trout Salmo trutta (Bernatchez, 2001; Bernatchez and Osinov, 1995), roaches *Rutilus* (Kotlík et al., 2008; Levin et al., 2017), and including Circum Black Sea barbels (Kotlík et al., 2004). During some periods usually associated with glaciation periods, when the level of Ocean dropped, the Black Sea level was lower by 100 (150) m (Yanina, 2014) and became a freshwater lake with a surface half of its current size. The last freshwater phase of the Black Sea terminated approximately 7150 years BP (Ryan et al., 1997). Despite the Black and Caspian Seas connecting several times during the Late Pleistocene sub-epoch via the Manych depression, no haplotypes of the Black Sea lineages were detected in the Caspian Sea basin.

We suggest that shallower divergence between members of the Black Sea group of species compared to such in the Caspian group is due to multiple repeated contacts between isolated lineages and sub-isolated populations during freshwater phases of the Black Sea water body. Therefore, glaciation deglaciation cycles have influenced the dispersal of circum Black Sea populations and could significantly upset speciation and divergence via a homogenizing effect on genetic structure. At the same time, an example of *B. rionicus*, with a well-diversified population structure in both mtDNA and nDNA loci that is well diverged from other Black Sea lineages and is geographically well restricted, provides evidence for local refugium probably existing for a long time during climatic oscillations.

### 4.4. Taxonomical implications and distributional notes

The most obvious taxonomic outputs of the present study are validation of the Rioni barbel *B. rionicus* and synonymization of the Sevan barbel *B. goktschaicus* with Kura barbel *B. cyri*. We confirmed revalidation that the *Barbus rionicus* was widely distributed in Western Transcaucasia, as inferred from our genetic data. The range of this species includes the Transcaucasian rivers from the Choruh basin northward to the Bzyb River, i.e., approximately 260 km along the Black Sea coast. Nevertheless, the northern range boundary of *B. rionicus* is permeable for the adjacent northern species, *B. escherichii* (loc. 42 in Fig. 1). Another species, *B. artvinica*, was described from the Choruh basin with *B. rionicus* simultaneously in the same book by Kamensky (1899); however, the Choruh population is genetically identical to other populations of *B. rionicus*. Therefore, both names are valid for the selection of only one of them, according to the International Code of Zoological Nomenclature (http://iczn.org/code). We accept the *B. rionicus* name as it has already been used (Bayçelebi et al., 2015).

No divergence between *B. goktschaicus* and *B. cyri* was detected by all studied markers. The Lake Sevan with tributaries being a range for *B. goktschaicus* was recently formed and isolated from the Araks basin approximately 20 000 years ago by the formation of an unpassable Varser waterfall on the Hrazdan R., which is the single outlet of Lake Sevan (Milanovskiy, 1957; Sarkissyan, 1962). Based on phenotypic data, Dadikyan (1986) recognized Sevan barbel as subspecies of the Kura barbel. At the same time, the Sevan population system is unique and composed of three sympatric forms (Chikova, 1955) under threat and protected by the list of the Red Data book of the Armenian Republic (Aghasyan and Kalashyan, 2010).

Our sampling covered type localities or type basins in the Kura system for some taxa that were described by Kamensky (1899), particularly *B. bortschalinicus* (loc. 23 in Fig. 1) and *B. toporovanicus* (loc. 19, 29 in Fig. 1). They were synonymized with *B. cyri* by Berg (1914). Our genetic data confirmed their identity with *B. cyri*.

The Crimean barbel *B. tauricus* was described from the Salgir R. by Kessler (1877). The Crimean populations are genetically heterogeneous and are similar to some populations of *B. escherichii* Steindachner, 1897, which are distributed around the Black Sea. The *B. tauricus* / *escherichii* complex of populations have a complicated history of migrations, re-colonizations of rivers discharged to Black Sea, and gene flow between lineages and populations (see Kotlík et al., 2004 and the present study). Another species, *B. oligolepis* Battalgil, 1941, which is known to be from the Marmara Sea basin, also belongs to the *B. tauricus* / *escherichii* complex, as inferred from the CÒI phylogeny (Fig. S1). At the same time, a couple of samples from the southern coast of the Black Sea, which were previously assigned as *B. escherichii*, are well diverged compared to other samples and might represent a different species (noted as *Barbus* sp. in the present study). Noticeably, there is another species complex, *Barbus petenyi* Heckel, 1852, among the barbels inhabiting the Danube basin, which also belonged to the Black Sea drainage(Antal et al., 2016; Kotlík et al., 2002; Žutinić et al., 2014).

## Acknowledgments

We are very thankful to O. Artaev, D. Karabanov, M. Matvejew, D. Palatov, A. Prokin, M. Saprykin, I. Turbanov, and A. Yakimov for their help in the field as well as to N. Bogutskaya for consultation on the taxonomy. The study was supported by Russian Science Foundation (grant no. 15-14-10020).

